# An ancient integration in a plant NLR is maintained as a *trans*-species polymorphism

**DOI:** 10.1101/239541

**Authors:** Helen J. Brabham, Inmaculada Hernández-Pinzón, Samuel Holden, Jennifer Lorang, Matthew J. Moscou

## Main text

Plant immune receptors are under constant selective pressure to maintain resistance to plant pathogens. Nucleotide-binding leucine-rich repeat (NLR) proteins are one class of cytoplasmic immune receptors whose genes commonly show signatures of adaptive evolution^1,2^. While it is known that balancing selection contributes to maintaining high intraspecific allelic diversity, the evolutionary mechanism that influences the transmission of alleles during speciation remains unclear. The barley *Mla* locus has over 30 described alleles conferring isolate-specific resistance to barley powdery mildew and contains three NLR families (*RGH1*, *RGH2*, and *RGH3*)^3^. We discovered (using sequence capture and RNAseq) the presence of a novel integrated Exo70 domain in RGH2 in the *Mla3* haplotype. Allelic variation across barley accessions includes presence/absence of the integrated domain in RGH2. Expanding our search to several Poaceae species, we found shared interspecific conservation in the RGH2-Exo70 integration. We hypothesise that balancing selection has maintained allelic variation at *Mla* as a *trans*-species polymorphism over 24 My, thus contributing to and preserving interspecific allelic diversity during speciation.

Recognition of plant pathogens is mediated by membrane and cytoplasmic immune receptors that recognise pathogen-derived molecules, including effectors^4^. Cytoplasmic immune receptors recognise secreted pathogen effectors directly, or indirectly through monitoring the molecular status of host proteins targeted by pathogen effectors^5^. At the population level, plant immunity to pathogens is conferred, in part, through the maintenance of diverse allelic variants at immune receptor loci. Pathogen pressure influences the relative frequency of resistance alleles in plant populations, with selection fluctuating due to pathogen evolution. Mutations that generate novel approaches to manipulate a host by immune suppression or nutrient acquisition, for example, are selected for in the pathogen. Similarly, novel forms of pathogen recognition are selected for in the plant. A recent area of intensive research in plant immunity includes the identification of integrated non-canonical domains in NLRs, which are thought to serve as baits for pathogen effectors, and thus, facilitate efficient immune responses^6–8^. It is hypothesised that integrated domains are released from purifying selection for host function, and are free to adapt specifically for pathogen recognition^6^.

*Blumeria graminis* f. sp. *hordei* (barley powdery mildew) is a pandemic disease of barley (*Hordeum vulgare*). A major source of isolate-specific resistance to *B. graminis* f. sp. *hordei* is found at the multi-allelic *Mla* locus, which contains the three NLR families *RGH1*, *RGH2*, and *RGH3*^3^. All known functional alleles of the *Mla* gene belong to the *RGH1* gene family^9^. Considerable haplotype variation has been identified based on sequencing *RGH1* genomic fragments, with alleles sharing >90% protein identity^3,9^. Specific residues of *Mla* proteins are under positive selection, particularly in the LRR region, which is known to determine the specificity of *B. graminis* f. sp. *hordei* recognition^9,10^. In the barley accession Morex, the *Mla* locus contains members of all three NLR families that are organised in three gene-rich regions interspersed with transposable elements^3^. In addition, a near identical 40-kb tandem duplication includes members from all three NLR families^3^. Morex contains *RGH1bcd*, an allele of *Mla* pseudogenised due to a retrotransposon insertion. We set out to understand the diversity of the *RGH* gene families across *Mla* haplotypes. To do so, we performed RNAseq on leaf tissue derived from diverse barley accessions, including elite, landrace, and wild barley. Sequence variation and presence/absence variation in expression of *RGH1*, *RGH2* and *RGH3* were identified across accessions (Fig. 1a). *RGH1* was the most prevalent of all *RGH* gene families with variation including intact open reading frames (ORF), pseudogenised genes, and in two cases, absence at the expression level. For all haplotypes sequenced, *RGH2* was always present with *RGH3*. Presence of *RGH3* without *RGH2* was seen in four accessions, and only in those also containing a pseudogenised *Mla* allele. Strikingly, we observed the presence of an integrated Exo70 domain within RGH2 in five haplotypes (accessions Baronesse, Duplex, Finniss, HOR 1428, and Maritime). Allelic variants of RGH2 in other accessions contain a coiled coil integrated domain derived from RGH1 or an in-frame full length RGH1 family member. Using sequence capture designed on the entire *Mla* locus and PacBio circular consensus sequencing^11^, the integrated *RGH2-Exo70* was confirmed in the barley cultivar Baronesse (Fig. 1c). Furthermore, *RGH2-Exo70* and *RGH3* in Baronesse were found in head-to-head orientation (Fig. 1b), and based on other similar structural examples we hypothesise they function as a pair^12^. Head-to-head orientation of *RGH2* and *RGH3* was also observed in other grass species (Fig. 1b).

**Fig. 1.**
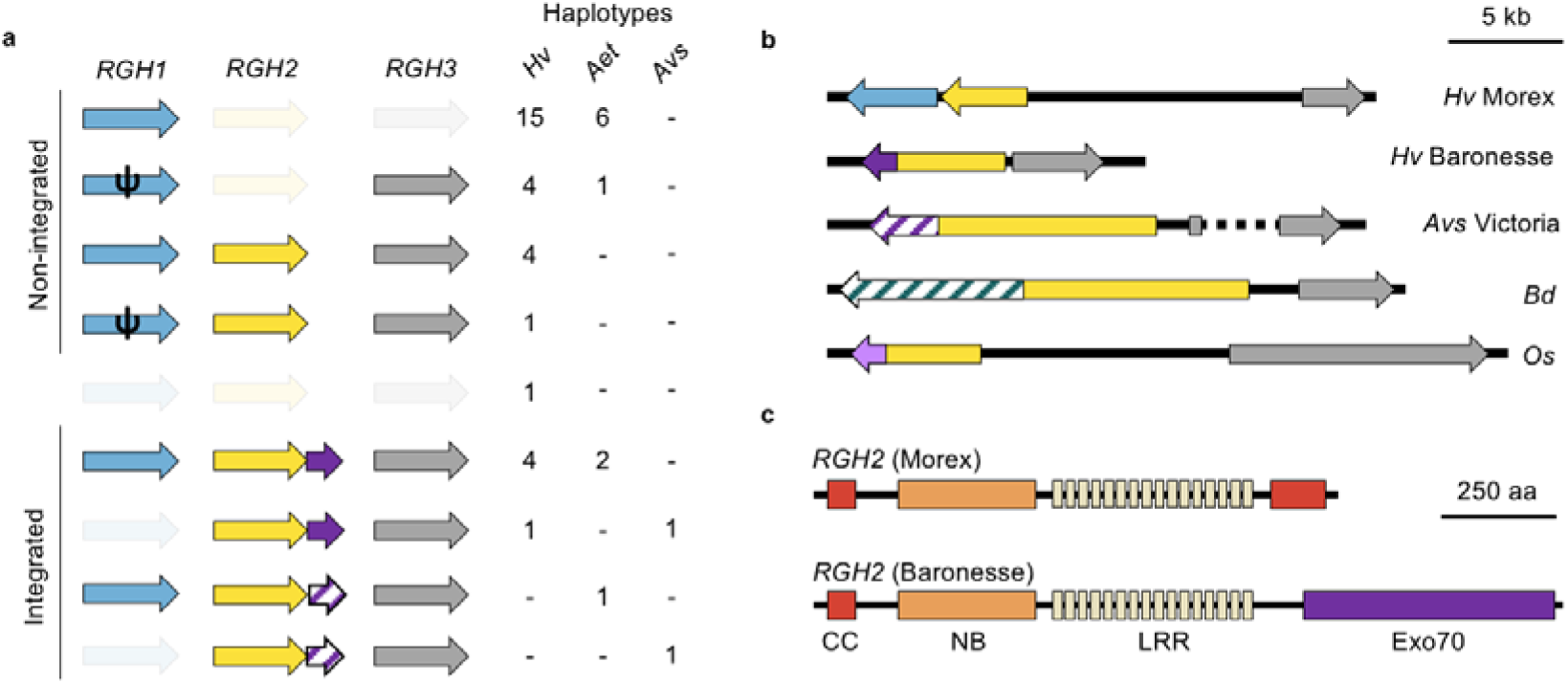
Intra- and inter-specific variation in *RGH1, RGH2*, and *RGH3*. RGH1, RGH2 and RGH3 family members are shown in coloured arrows (blue, yellow, and grey, respectively). (a) Haplotype variation in leaf expression of *RGH1*, *RGH2*, and *RGH3* within barley (*Hordeum vulgare*; *Hv*), *Aegilops tauschii* (*Aet*), and oat (*Avena sativa*; *Avs*). Presence of a complete coding sequence is shown in bold arrows, absence with faded arrows, pseudogenised *RGH1* with the symbol ψ, integration of Exo70 with purple arrows, and out-of-frame integrated Exo70 with hatched shaded arrows. Haplotypes is defined as the number of haplotypes containing different gene combinations. (b) Genomic structure of the region encompassing *RGH2* and *RGH3* across Poaceae species showing conserved head-to-head orientation of *RGH2* and *RGH3*. Variation is observed in RGH2 integrated domains with barley accessions Morex (*mla*)^3^ and Baronesse (*Mla3*/*Rmo1*)^9,30^ containing *RGH2*-*Exo70* (yellow-purple arrow), oat accession Victoria containing *RGH2-Exo70* (yellow-purple arrow), *Brachypodium distachyon* (*Bd*) containing *RGH2-Receptor-like-kinase* (yellow-teal arrow), and rice (*Oryza sativa*; *Os*) containing *RGH2-Thioredoxin* (yellow-light purple arrow). *A. sativa* accession Victoria and *B. distachyon* integrated *Exo70* are out-of-frame with *RGH2*, indicated with the hatched shaded arrows. (c) Protein model of non-integrated RGH2 and integrated RGH2-Exo70. Individual domains include coiled coil (CC; red), nucleotide-binding (NB; orange), leucine-rich repeats (LRR; cream), and Exo70 (purple).

Exo70 is one of the eight subunits comprising the exocyst complex, involved in the tethering of post-Golgi secretory vesicles to the plasma membrane prior to SNARE complex-mediated membrane fusion^13–15^. In plant genomes, the core exocyst subunits of Sec3, Sec5, Sec6, Sec8 and Sec10 are retained in few or single copies, whereas Exo70 has experienced dramatic proliferation into multiple gene families^16^. Diversification is characteristic of sub-functionalisation or neo-functionalisation^17^ and Exo70 genes are involved in diverse roles in morphogenesis^15^, development^18^, and immunity^19^. Cvrcóková *et al*. (2012) comprehensively described members of the exocyst complex throughout land plants using representative species from both monocots and dicots^16^. We annotated the *Exo70* gene families from several sequenced grass species including *Brachypodium distachyon*, barley, *Oryza sativa* (rice), *Oropetium thomaeum*, *Sorghum bicolor*, *Setaria italica*, and *Zea mays* (maize). The *Exo70F* and *Exo70FX* clades were found to be greatly expanded within these grasses (Fig. 2a). Using maximum likelihood phylogenetic analysis, we discovered that the integrated *Exo70* originated from *Exo70F1* (Fig. 2b). Approximately 87% of the coding sequence of *Exo70F1* is integrated in *RGH2*. Members of *Exo70F* and *Exo70FX* gene families are known to have a role in immunity. Barley *EXO70F*-*like* (*ExoFX11b.a*) is essential for full penetration resistance to *B. graminis* f. sp. *hordei*^20^. Rice Exo70F3 binds the *M. oryzae* effector AVR-Pii, and this interaction is essential for *Pii* (NLR) mediated resistance^21,22^. The integration of Exo70F1 in RGH2 suggests a potential role in immunity either through localisation of the NLR or as a decoy for effector recognition.

**Fig. 2.**
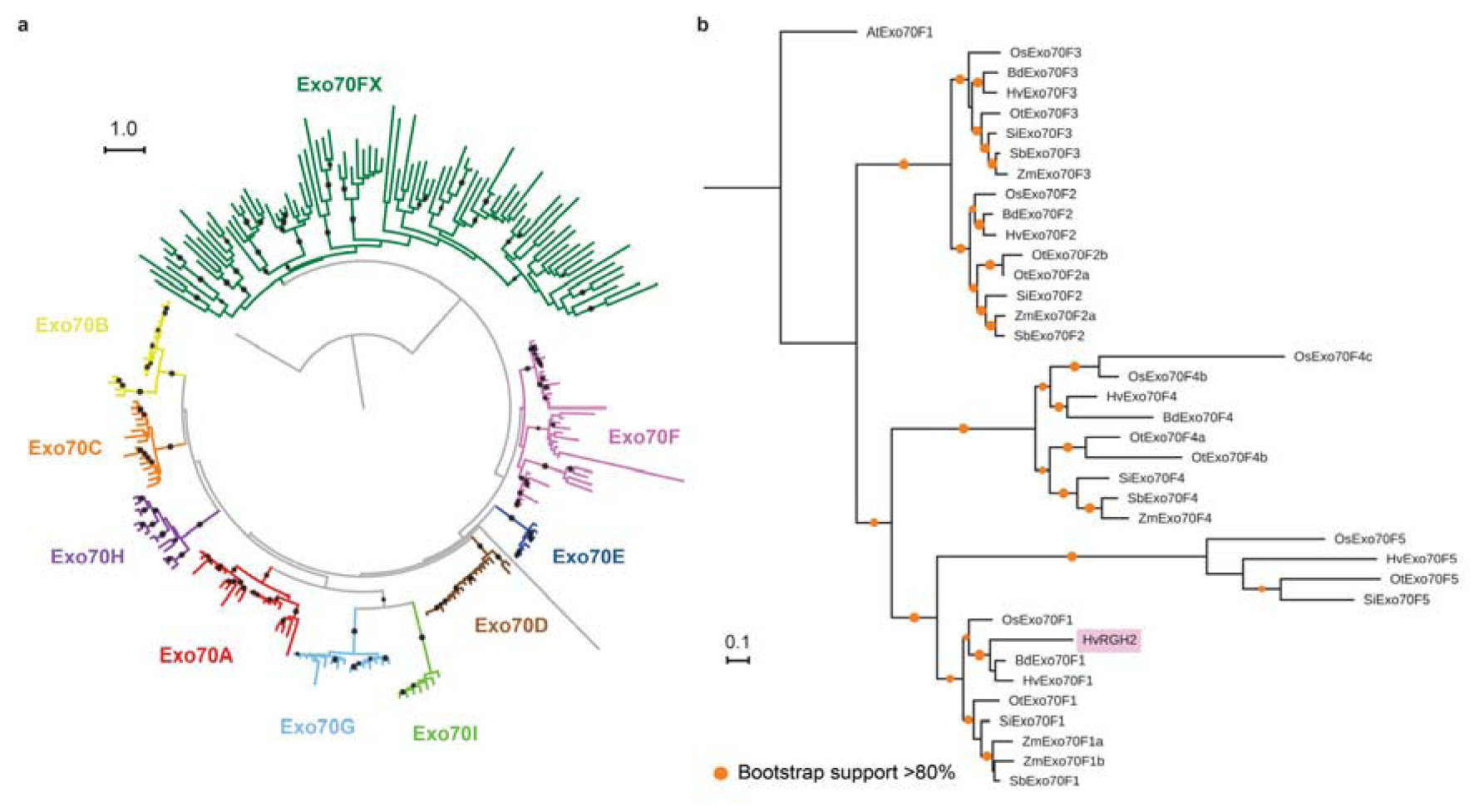
*Exo70F1* is the donor of the *Exo70* integration in *RGH2*. (a) Maximum likelihood phylogenetic tree of Exo70 proteins from seven grass species (see below; Supplementary Data 1), including barley *RGH2* integrated *Exo70* denoted HvRGH2. *Saccharomyces cerevisiae* Exo70 gene (YJL085W) was used as an outgroup. Black dots represent bootstrap support greater than 80% based on 1,000 bootstraps. (b) Maximum likelihood phylogenetic tree of Exo70F proteins from seven grass species (see below). The origin of the integrated Exo70 in RGH2 is Exo70F1 family (highlighted in pink). *Arabidopsis thaliana* Exo70 gene (AT5G50380; AtExo70F1) was used as an outgroup. Orange dots represent bootstrap support greater than 80% based on 1,000 bootstraps. Species included in the analysis were: *B. distachyon* (*Bd*), barley (*Hv*), rice (*Os*), *Oropetium thomaeum* (*Ot*), *Sorghum bicolor* (*Sb*), *Setaria italica* (*Si*), maize (*Zea mays*; *Zm*).

Integration of *Exo70F1* in *RGH2* was previously observed in the D genome of wheat (*Triticum aestivum* L.) in cloning *Sr33* on chromosome 1D^23^. Presence of integrated *Exo70F1* in wheat and barley prompted us to investigate its conservation across diverse grasses to establish its potential as a *trans*-species polymorphism^2^. First, to understand whether allelic variation exists in *Aegilops tauschii* (the donor of the D genome), we analysed several publically available leaf transcriptomes^24^. We observed allelic variation in the presence or absence of *RGH2* and *RGH3* expression, with *RGH2* present as the integrated *Exo70F1* allele (Fig. 1a). We did not identify an allele of *RGH2* without the *Exo70F1* integration. We expanded our search to include additional species outside the Triticeae with sequenced genomes or publically available leaf RNAseq data. Sequence analysis found allelic variation and interspecific conservation in the *RGH2-Exo70F1* integration across diverse Poaceae species. In oat, (*Avena sativa*) *RGH2* and *RGH3* were present in two accessions, whereas *RGH1* was absent (Fig. 1a). The integrated *Exo70F1* was in frame in the accession Kanota, whereas a single base pair InDel causes an out-of-frame *Exo70F1* in Victoria. Integration of *Exo70F1* in *RGH2* in the genome was confirmed using sequence capture and PacBio sequencing of the oat accession Victoria. Using maximum likelihood phylogenetic analysis, we found that integrated *Exo70F1* derived from Triticeae species, *A. sativa*, and *Holcus lanatus* form a distinct clade from non-integrated *Exo70F1* (Fig. 3). *B. distachyon* forms the outgroup of this integrated clade, suggesting that integration of *Exo70F1* in *RGH2* occurred after speciation of *B. distachyon* but prior to radiation of the Poeae and Triticeae. The *RGH2* ortholog in *B*. *distachyon* encodes an NLR with a C-terminal integrated receptor-like kinase with intact transmembrane and extracellular lectin domains. *RGH2* orthologs in oat and the *Ae. tauschii*-derived D genome of wheat contain integrated *Exo70F1* that is not in frame, suggesting that gene exchange through unequal crossovers, gene conversion, or mutation can interrupt the ORF. The differentiation of integrated and non-integrated *Exo70F1* after *B*. *distachyon* speciation, but prior to Poeae-Triticeae radiation dates this integration at 24 Mya (CI: 18.4 to 29.8 Mya)^25^, and is indicative of a *trans*-species polymorphism.

**Fig. 3.**
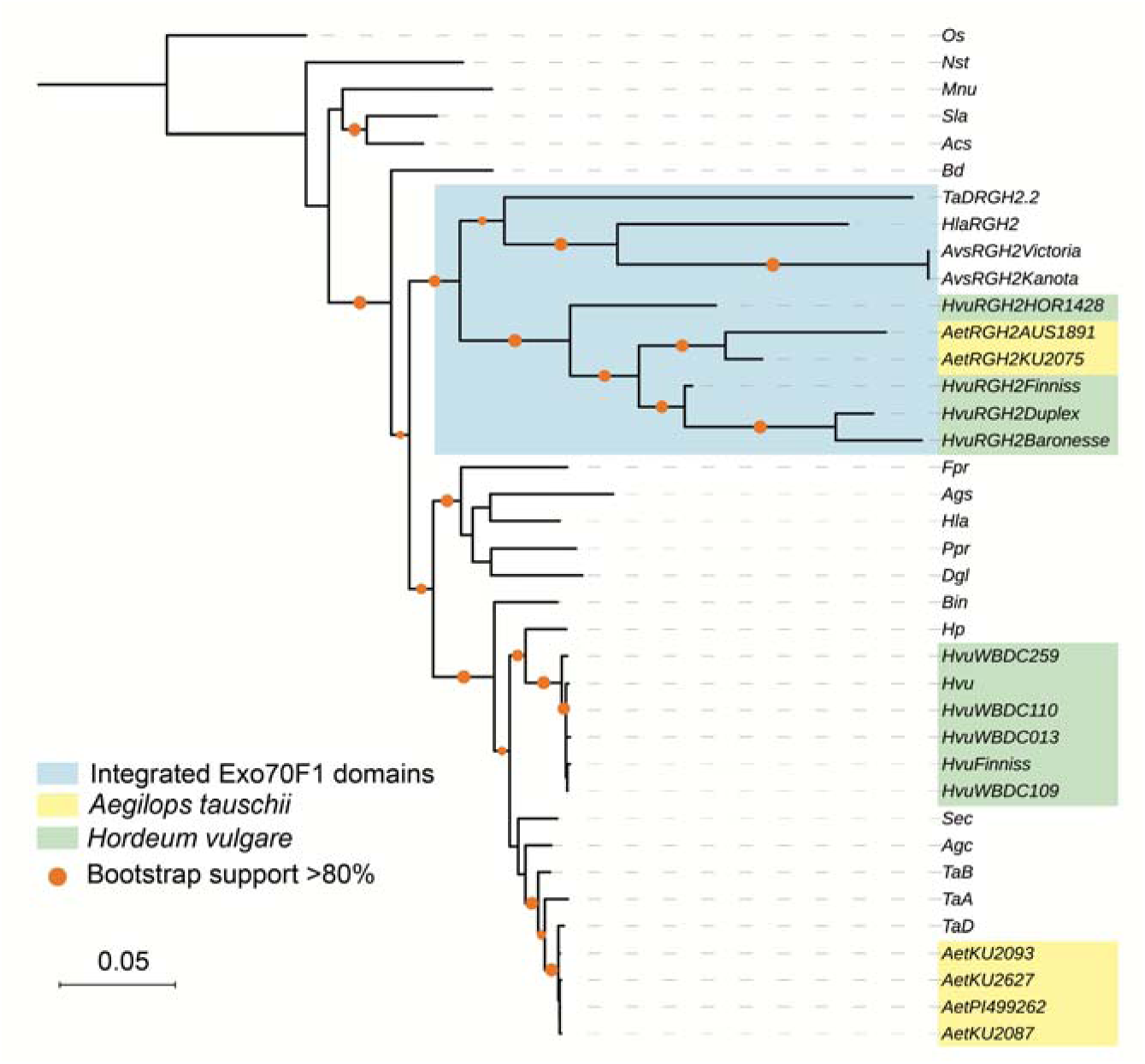
*Exo70F1* integration in *RGH2* occurred prior to the Poeae*-*Triticeae radiation. Maximum likelihood phylogenetic tree was performed on codon aligned *Exo70F1* genes from 19 grass species (Supplementary Data 1). Integrated *Exo70F1* highlighted in blue. Branch support was generated using 1,000 bootstraps, with orange dots designating support greater than 80%. Rice *Exo70F1* (*OsExo70F1*; *Os01g69230.1*) was used as an outgroup. Scale bar shows nucleotide substitutions per site. Species included in the analysis were: *Achnatherum splendens* (*Acs*), *Ae. tauschii* (*Aet*), *Agropyron cristatum* (*Agc*), *Agrostis stolonifera* (*Ags*), *A. sativa* (*Avs*), *B. distachyon* (*Bd*), *Bromus inermis* (*Bin*), *Dactylis glomerata* (*Dgl*), *Festuca pratensis* (*Fpr*), *Holcus lanatus* (*Hla*), *H. pubiflorum* (*Hp*), barley (*Hvu*), rice (*Os*), *Melica nutans* (*Mnu*), *Nardus stricta* (*Nst*), *Poa pratensis* (*Ppr*), *Secale cereale* (*Sec*), *Stipa lagascae* (*Sla*), *Triticum aestivum* (*TaA*, *TaB*, *TaD* subgenomes), maize (*Z. mays*; *Zm*). Clusters were formed due to 100% identical sequence and are listed in Supplementary Table 1.

We found that all grass *Exo70* gene families are under strong purifying selection (Supplementary Table 2). We hypothesise that integration of *Exo70F1* into *RGH2*, and its role as a proposed ‘decoy’ domain, would relax purifying selection compared to non-integrated *Exo70F1*. To understand the impact on the molecular evolution across *Exo70F1*, we performed branch-specific tests for variable levels of selective pressure. Evidence of relaxed purifying selection was observed for the integrated *Exo70F1* clade (ω_a_ = 0.423), compared to stronger purifying selection for non-integrated domains of *Exo70F1* (ω_0_ = 0.102) (Supplementary Table 3, Supplementary Fig. 1). Evidence of relaxation confirms the release of integrated domains from strong purifying selection that preserves the endogenous activity of their non-integrated counterparts^26^.

We were interested if, following incorporation, integrated *Exo70F1* has altered the evolution of *RGH2*. To investigate the potential co-evolution of *RGH2* and the integrated *Exo70F1*, we compared the phylogenies of *RGH1*, *RGH2*, and *RGH3* gene families; and *Exo70F1*. All *RGH* gene families predominantly exhibit species specific grouping (Supplementary Fig. 2, 3, 4). Variation is seen in the domain structure of RGH2 homologs; the majority contain an Exo70F1 integration however other domains such as coiled coil, receptor-like kinase (with lectin domains), thioredoxin and protein kinase are observed in orthologs (Supplementary Fig. 3). *RGH2* belongs to the NLR clade MIC1 comprising members known to contain diverse integrated domains^27^. Integrated *Exo70F1* have a distinct evolutionary history compared to their non-integrated counterparts. The integrated *Exo70F1* clade follows the species phylogeny, with the exception of *HvuRGH2HOR1428* and *TaDRGH2.2* from the reference genome of wheat (Fig. 3). Within the integrated clade two subclades are supported: one clade containing the *Aegilops* and barley alleles, and the other comprising wheat, *Phalaris arundinacea, Holcus lanatus*, and oat (Fig. 3). In contrast, *RGH2-Exo70F1* and non-integrated *RGH2* alleles do not show the same clade distinction; therefore, integrated status has not altered *RGH2* evolutionary history and co-evolution has not occurred (Supplementary Fig. 3).

Until now, it was unclear how selective forces in the plant-pathogen interaction influence the preservation of diverse alleles through plant speciation events because previous evolutionary analyses were limited by high rates of intraspecific and interspecific variation of NLRs. We used the integrated *Exo70F1* domain within *RGH2* as an evolutionary footprint to track the history of the *Mla* locus through speciation of the grasses. We observed presence/absence variation in *RGH2*-*Exo70F1* across barley and presence of *RGH2*-*Exo70F1* alleles in five grass species, indicating the passage and retention of ancestral alleles through speciation. This establishes a *trans*-species polymorphism originating 24 Mya^25^, and maintained to present day (Fig. 4). Long-lasting *trans*-species polymorphisms are maintained through balancing selection, as in its absence, mutations within a population are either lost, achieve fixation, or exist in a frequency-dependent manner^2^. Previous work in plant immunity established time scales of long-term maintenance of polymorphic sites in stress response genes between *A. thaliana* and *Capsella rubella*^28^ over 5 Mya, and presence/absence polymorphisms in *RPS5* in *A. thaliana* due to long-term balancing selection^29^ over 2 Mya. Using genomes and transcriptomes from species across the Poaceae, we have unravelled the evolutionary history of a shared integrated NLR within the grasses, and shown that allelic diversity is present as a *trans*-species polymorphism at *Mla*. The association of the *Mla* locus with multiple pathogen recognition^30^ emphasises the impact of diverse pathogen populations on the evolutionary forces shaping immune receptor diversity in the Poaceae.

**Fig. 4.**
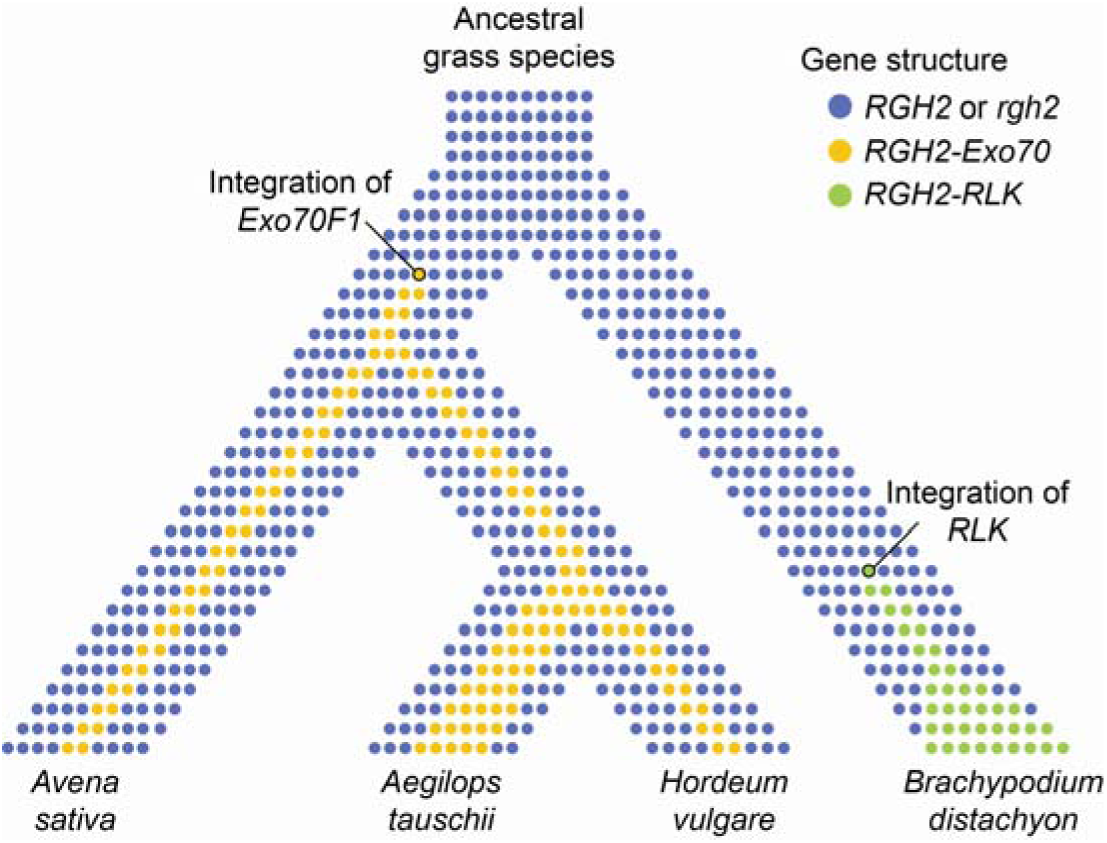
An evolutionary model of integrated and non-integrated *RGH2* transmission during the speciation of Poeae and Triticeae. Horizontal lines of dots represent the gene pool across a species at a given point in time. Branch points represent speciation events. Coloured dots represent allelic variants of *RGH2*; blue dots represent non-integrated alleles, yellow dots represent *RGH2-Exo70* integrations, and green dots show *RGH2-Receptor-like kinase* (*RGH2-RLK*) integration in *B. distachyon*. Following speciation of *B*. *distachyon*, an allele of *RGH2* acquires the integrated *Exo70F1*. Integrated and non-integrated alleles of *RGH2* are maintained within the population though balancing selection, and are sustained through the speciation of *Avena* and other Poeae species, and subsequently the Triticeae. Long-term balancing selection preserves the *RGH2* allelic pool as a *trans*-species polymorphism.

## Methods

### Plant material

A full inventory of plant species and accessions used in this study are available in Supplementary Data 1.

### *RNAseq and* de novo *assembly*

First and second leaf tissue was harvested at 10 days after sowing of barley and oat accessions grown in the greenhouse. Tissue was flash frozen in liquid nitrogen and stored at −80 °C. Tissue were homogenized into a fine powder in liquid nitrogen-chilled pestle and mortars. RNA was extracted, purified, and quality assessed as described by Dawson *et al*. (2016)^31^. RNA libraries were constructed using Illumina TruSeq RNA library preparation (Illumina; RS-122-2001). Barcoded libraries were sequenced using either 100 or 150 bp paired-end reads. Library preparation and sequencing was performed at either the Earlham Institute (Norwich, United Kingdom) or BGI (Shenzhen, China). Quality of all RNAseq data was assessed using FastQC^32^ (0.11.5). Trinity^33^ (2.4.0) was used to assemble *de novo* transcriptomes using default parameters and Trimmomatic^34^ for read trimming. *Exo70F1*, *RGH1*, *RGH2*, and *RGH3* were identified in *de novo* assemblies using BLAST+ (v2.2.9)^35^.

### Development of the barley and oat sequence capture

The barley capture library TSLMMHV1 is composed of 99,421 100 mer baits with 2x coverage over the target space. Targeted sequences include repeat masked *Mla* locus from Morex^3^, all cloned alleles of *Mla^9,10,36-38^*, the *Mlo* locus^39^, the *Rpg1* locus^40^, and the *rpg4*/*Rpg5* locus^41^. In addition, the capture library targets the barley NLR gene space identified in genomic sequence from barley accessions Barke, Bowman, and Morex, full length cDNA derived from barley accession Haruna Nijo, and transcriptomes of barley accessions Abed Binder 12, Baronesse, CI 16153, CIho 4196, Manchuria, Pallas, Russell, and SusPtrit. The capture library TSLMMAS1 is composed of 29,672 100 mer baits with 2x coverage over the target space. Targeted sequences include NLR containing contigs from *de novo* assembled transcriptomes of oat accessions Kanota and Victoria.

For both libraries, NLR gene space was identified using the following approach. For transcriptomes, TransDecoder^42^ (v4.1.0) was used to identify and translate ORFs. Using a similar strategy as Jupe *et al*. (2012)^43^, we developed a motif set using MEME^44^ trained on a random randomized proportional sample of NLR^45,46^ from rice (N=35) and *B. distachyon* (N=17). The MEME motifs spanned the CC domain (motifs 4, 11, 13, and 15), NB domain (motifs 1, 2, 3, 5, 6, 7, 8, 10, 12, and 14), and the LRR domain (motifs 19, 9, 20, 16, 17, and 18) (Supplementary Data 3). All the identified motifs in the NB are similar to those previously defined by Meyers *et al*. (2003)^47^ in *A. thaliana*. A MAST significance threshold of 1e-20 was selected based on its ability to identify all annotated NLRs within *B. distachyon* and rice. For barley whole genome shotgun assemblies, all six ORFs were translated for each contig and concatenated into a single peptide sequence for the forward and reverse strand. Translated genomic contigs were scanned using FIMO, which assesses all twenty MEME-generated motifs independently. Contigs were included in the capture if one of two conditions were met: (1) at least one CC and two NB motifs were present or (2) at least two NB and one LRR motifs are present in the translated sequence strand.

Next, redundancy within the sequence capture template was removed. We fragmented the input data set into 100 bp fragments with a scanning window of 25 bp and performed BLASTn onto the entire data set. Any sequence found to have identity of 95% or higher was considered redundant other than the original site. The first occurrence of the sequence would be retained and the others were masked. While this approach removes redundancy in the data set, the inclusion of extensive genomic sequence will introduce repetitive sequence that can produce competition in the sequence capture due to the high copy number of repetitive sequence in the barley genome. Therefore, two approaches were used to remove repetitive sequence in the sequence capture design. All loci were repeat masked based using RepeatMasker^48^ (v4.0.5) using default and Triticeae-specific repeat databases. As repeat databases are not complete, we applied genomic masking of the capture design. To do so, we fragmented the input data set into 100 bp fragments with a scanning window of 50 bp and performed BLAST onto the barley Morex WGS assembly. A threshold of eight or fewer copies was found selected to balance between copy number variation within NLRs and avoiding the inclusion of repetitive sequence.

### DNA extraction and sequencing library preparation

Total genomic DNA was extracted from leaf tissue according to a CTAB method^49^. In brief, 3 g of leaf tissue were ground on liquid N_2_ and homogenized with 20 mL of CTAB extraction buffer (2% CTAB, 100 mM Tris-HCl pH8.0, 20 mM EDTA pH8.0, 1.4 M NaCl, 1% β-Mercaptoethanol). Samples were incubated for 30 min at 65°C followed by two chloroform extractions and ethanol precipitation. DNA was then resuspended in 1x TE, 50 μg/mL RNase A solution and incubated for 1 h at 37°C. DNA was subsequently precipitated with 2.5 volumes of ice-cold 95% ethanol and resuspended in 1x TE. Quantification of DNA samples was performed using a Nanodrop spectrophotometer (Thermo Scientific) and the Qubit dsDNA HS Assay Kit (Molecular Probes, Life Technologies). DNA samples were normalized to 3 µg and sheared to an average length of 3-4 kb using a Covaris S2 sonicator with the following settings: Duty Cycle 20%, Intensity 1, Cycle Burst 1000, Time 600 s, Sample volume 200 µL. After sonication, a small aliquot was assayed by gel electrophoresis and additional size selection was carried out using 0.4x Agencourt AMPure XP beads (Beckman Coulter Genomics). Samples were then end repaired followed by 3’dA addition using the NEBNext Ultra DNA Library Prep Kit for Illumina (New England Biolabs). Illumina sequencing adapters were ligated onto the ends and following purification with AMPure XP beads, the DNA was PCR amplified (8 cycles) using indexed PCR primers (NEBNext Multiplex Oligos for Illumina, New England Biolabs) and the Illumina PE1.0 PCR primer. After purification using AMPure XP beads, quality assays were performed with a Bioanalyzer DNA 1000 chip (Agilent) and the Qubit dsDNA assay to determine the average fragment sizes and concentrations.

### Target enrichment and sequencing

Enrichment and sequencing was carried out as described by Witek *et al*. (2016)^11^. Briefly, DNA sequencing libraries was enriched according to the MYbaits protocol (MYbaits User Manual version 2.3.1) and using MYbaits reagents (MYcroarray). Briefly, 500 ng of the prepped library was hybridized in hybridization buffer (10x SSPE, 10X Denhardt’s solution, 10 mM EDTA, 0.2% SDS) to the biotinylated RNA baits for 20 h at 65°C on a thermocycler. After hybridization bound DNA was recovered using magnetic streptavidin-coated beads as follows: the hybridization mix was added to 30 µL Dynabeads MyOne Streptavidin C1 (Invitrogen, Life Technologies) that had been washed 3 times and resuspended in binding buffer (1 M NaCl; 10 mM Tris-HCl, pH 7.5; 1 mM EDTA). After 30 m at 65°C, beads were pulled down and washed three times at 65°C for 10 m with 0.02% SSC/0.1% SDS followed by resuspension in 30 µL of nuclease-free water. Library was then PCR amplified (26 cycles) using Kapa HiFi HotStart Ready Mix (Kapa Biosystems) and Illumina P5 and P7 primers. The amplified library was size fractionated with the Sage Scientific Electrophoretic Lateral Fractionator (SageELF, Sage Science) using a 0.75% SageELF agarose gel cassette. Fractions with size distribution between 3 and 4 kb were pooled and purified with AMPure PB beads (Pacific Biosciences). Then, library was assembled for PacBio sequencing using the SMRTbell Template Prep Kit 1.0 (Pacific Biosciences) according the 2-kb Template Preparation and Sequencing protocol (www.pacificbiosciences.com/support/pubmap/documentation.html). PacBio RSII sequencing using C4-P6 chemistry was performed at the Earlham Institute (Norwich, UK), using four SMRT cells for each barley accession Baronesse and oat accession Victoria.

### PacBio assembly

PacBio circular consensus reads with at least three passes were used for genome assembly. Reads were trimmed to remove the adapter sequence (first and last 70 bp) and size selected to reads less than 4kb. We used Geneious^50^ (v10.2.3) De Novo Assembly with the following Custom Sensitivity parameters for assembly: don't merge variants with coverage over approximately 6, merge homopolymer variants, allow gaps up to a maximum of 15% gaps per read, word length of 14, minimum overlap of 250 bp, ignore words repeated more than 200 times, 5% maximum mismatches per read, maximum gap size of 2, minimum overlap identity of 90%, index word length 12, reanalyze threshold of 8, and maximum ambiguity of 4.

### Phylogenetic analyses

Exo70 coding and protein sequences were accessed from Department of Energy-Joint Genome Institute Phytozome database (https://phytozome.jgi.doe.gov), *A. thaliana* gene sequence was accessed from TAIR (www.arabidopsis.org), and barley *Exo70* gene families from the recently updated 2017 genome^51^. Substantial curation of the barley *Exo70* gene family was required and incorporated gene models from the 2012 genome^52^. Outgroups for complete Exo70, Exo70F, and *Exo70F1* phylogenetic trees were *Saccharomyces cerevisiae* Exo70 protein (YJL085W), *A. thaliana* Exo70F1 protein (AT5G50380), and *O. sativa Exo70F1* (*Os01g69230.1*), respectively. MUSCLE^53^ (v3.8.31) and PRANK^54^ (v.140603) were used for protein and codon-based sequence alignment using default parameters, respectively. Curation of the multiple sequence alignment for complete Exo70 and Exo70F gene family was used to remove sequences with less than 40% of the breadth of the alignment and to remove positions with more than 60% missing data. Gene families were identified based on bootstrap support in the phylogenetic tree (Fig. 2a), incorporating a previous annotation performed on the *Exo70* gene family^16^ (Supplementary Data 4). We required that 90% sequence coverage for inclusion in the *Exo70F1* phylogenetic tree. RAxML^55^ (v8.2.9) was used for phylogenetic tree construction using the PROTGAMMAJTT and GTRCAT models for protein and coding sequence alignment, respectively. Bootstrap support was determined for all phylogenetic trees, using a convergence test to confirm sufficient sampling.

*Exo70F1* homologs were identified from diverse Poales species^56^ using BLAST+ (Supplementary Data 1). The only species found without a non-integrated *Exo70F1* was *H. lanatus*. Alignment of *H. lanatus* RNAseq reads to *Dactylis glomerata Exo70F1* and *de novo* assembly using Geneious was used to reconstruct *H. lanatus Exo70F1*. Multiple sequence alignment and phylogenetic analysis of *Exo70F* gene families was used to establish *Exo70F1* orthology (Supplementary Fig. 5). The site of sequence conservation between integrated and non-integrated *Exo70F1* was determined based on codon-based sequence alignment.

### Molecular evolutionary analyses

Molecular evolutionary analyses were performed with PAML^57^ (v4.8) *codeml*. The F3x4 codon frequency model was used for all analyses. Codon-based sequence alignment using PRANK, phylogenetic analysis using RAxML, and codeml were used to estimate ω (d_N_/d_S_) for all Exo70 gene families (Supplementary Table 2). For molecular evolutionary analyses of the non-integrated and integrated *Exo70F1* gene family, four hypotheses were tested: H_0_: a single rate of ω_0_ across the entire tree, H_1_: a different rate of ω_1_ for the initial branch of the integrated *Exo70F1*, H_2_: a different rate of ω_a_ for integrated *Exo70F1* compared to non-integrated *Exo70F1* (ω_0_), and H_3_: three different rates of ω: ω_0_, non-integrated *Exo70F1* outside of the Poeae and Triticeae, ω_a_, integrated Exo70F1, and ω_b_, non-integrated *Exo70F1* from the Poeae and Triticeae (Supplementary Fig. 1, Supplementary Table 3).

### Data

All high-throughput sequencing data, *de novo* transcriptome assemblies, and *de novo* assembly of the NLR gene space of barley and oat are deposited in the NCBI BioProject PRJNA378334, PRJNA378723, PRJNA422803, and PRJNA422986. *De novo* transcriptome assemblies for publically available RNAseq data, multiple sequence alignments, and phylogenetic trees in Newick format are available from figshare (https://figshare.com). Scripts, analysis pipeline, and details associated with *Exo70* gene family curation can be found on the GitHub repository https://github.com/matthewmoscou/Exo70 (v1.0).

## Acknowledgements

The authors thank Tom Wolpert for excellent advice, enthusiasm, and curiosity, Phon Green for management of plant materials, William Jackson for the development of initial bioinformatic approaches for assembly of NLR gene space, Kamil Witek for advice on sequence capture and PacBio library preparation, Ksenia Krasileva for discussion on NLR-ID evolution, Simon Griffiths and Sophien Kamoun for their critical input, and Ryohei Terauchi and Hiromasa Saitoh for their open dialogue of science. This work was funded using core funding from the Gatsby Foundation and a BBSRC Institute Strategic Programme (BB/P012574/1). Additional support from the Perry Foundation was provided as a PhD studentship to HJB. JL was funded by grant 2016-67013-24736 from the USDA National Institute of Food and Agriculture.

## Author contributions

Conceived and designed the experiments: H.J.B., J.L., and M.J.M. Performed transcriptome sequencing and sequence capture: I.H-P. Performed bioinformatic analyses: H.J.B., S.H., and M.J.M. Analysed the data: H.J.B., S.H., J.L., and M.J.M. Wrote the paper: H.J.B. and M.J.M.

## Supplementary Information

**Supplementary Fig. 1.**
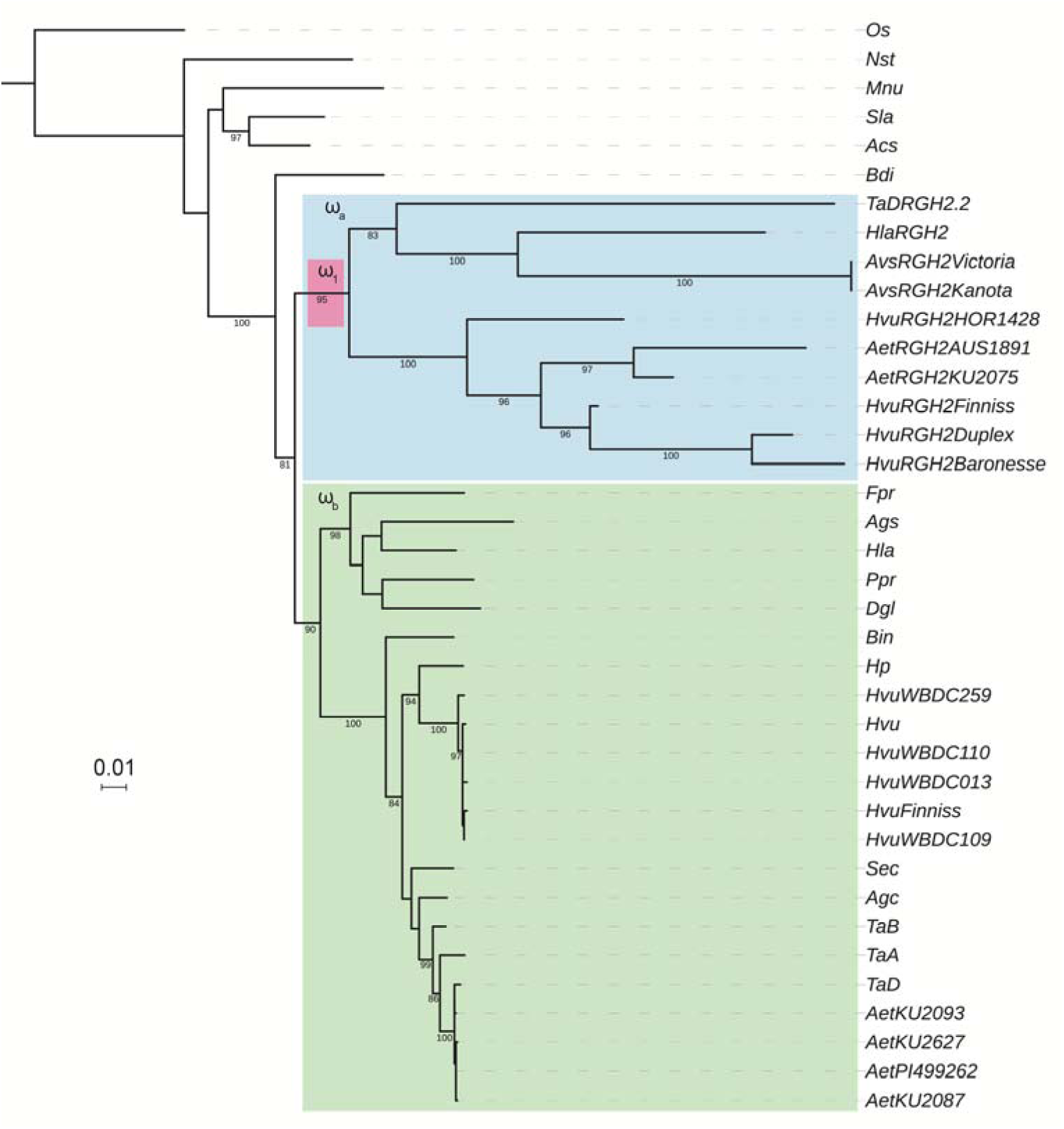
Exo70F1 maximum likelihood phylogenetic tree used for molecular evolutionary analyses. Branch and clade-based estimation of ω (d_N_/d_S_) are highlighted: ω_1_ in pink, ω_a_ in blue, and ω_b_ in green (Supplementary Table 3). Unit of distance is nucleotide substitutions per evaluated sites. Species abbreviations listed in Fig. 3. *HvuRGH2Maritime* clusters with *HvuRGH2Baronesse* due to 100% similarity. Rice (*Os*; *LOC_Os01g69230.1*) used as an outgroup.

**Supplementary Fig. 2.**
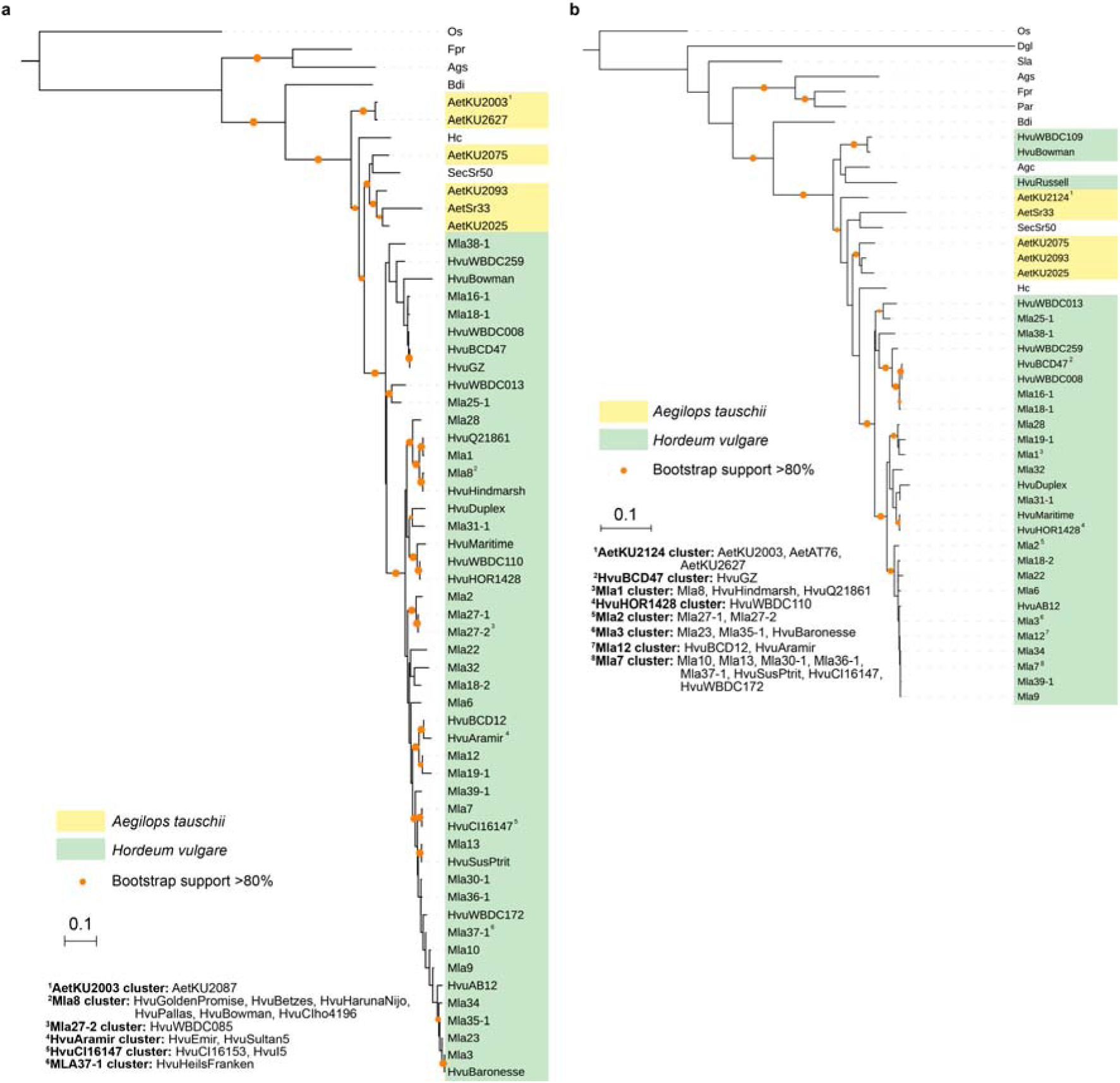
Maximum likelihood phylogenetic tree of *RGH1* homologs across the Poaceae. Alignments and phylogenetic trees of full length (a) and NB domain analyses (b) were performed. Clusters of identical sequence are indicated by superscript numbers and indicated in the legend. Branch support was generated using 1,000 bootstraps for both phylogenetic trees, with orange dots designating support from 80-100%. Species abbreviations listed in Supplementary Data 1. Rice (*Os*; *LOC_Os11g43700.1*) used as an outgroup.

**Supplementary Fig. 3.**
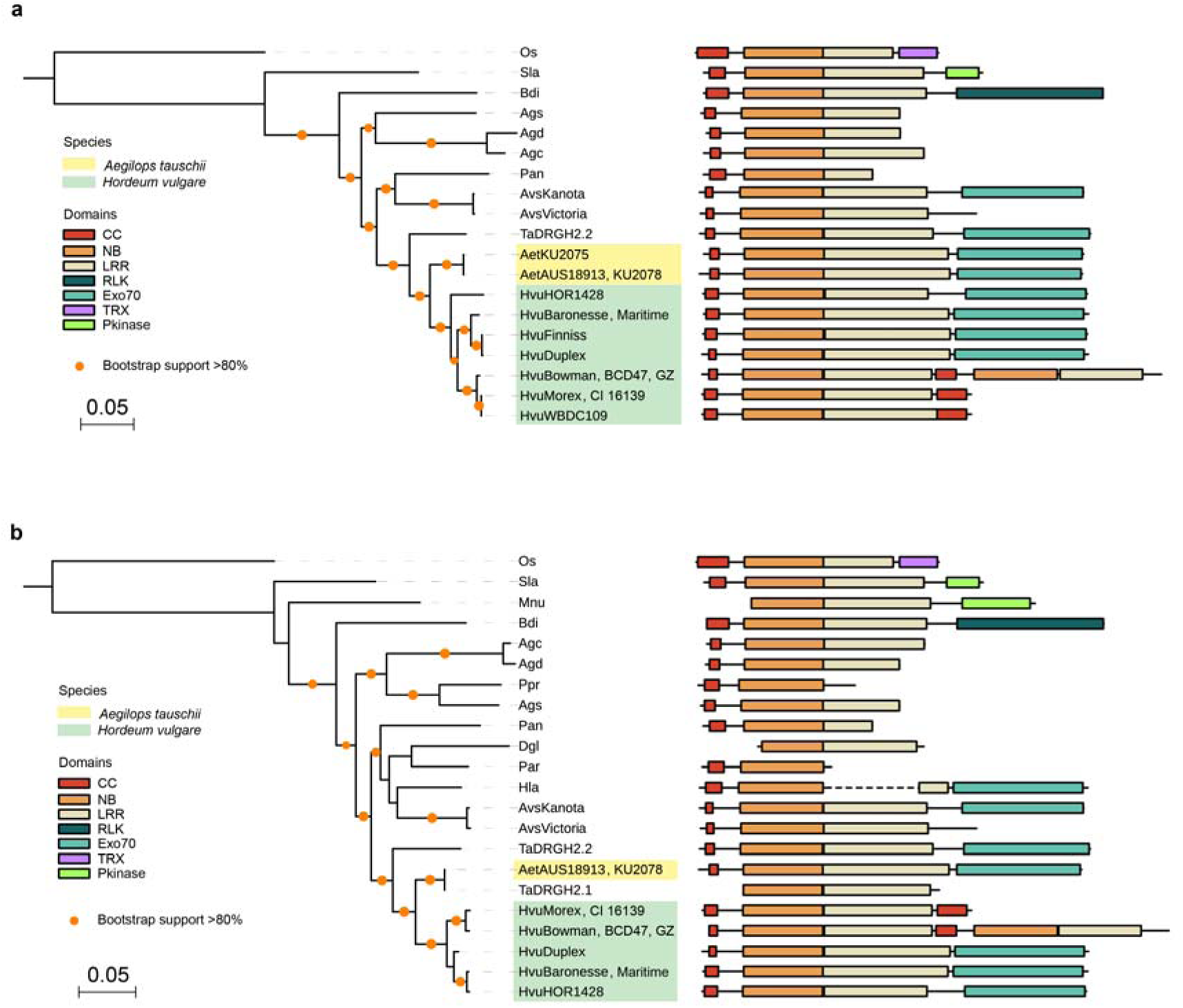
Maximum likelihood phylogenetic tree and domain structure of *RGH2* homologs across the Poaceae. Alignments and phylogenetic trees of full length (a) and NB domain analyses (b) were performed. *RGH2* domain structure represented on the right. RGH2 from *Ae. tauschii* and *H. vulgare* indicated with yellow and green, respectively. AvsVictoria integrated *Exo70F1* domain absent from domain structure due to frame-shift relative to RGH2. In addition to canonical domains (CC; red, NB; orange, LRR; cream), additional C-terminal integrated domains are observed in *RGH2* alleles and include thioredoxin (TRX; purple), protein kinase (Pkinase; light green), lectin (teal) and Exo70 (cyan). Dashed lines represent fragmented ORFs (only observed for *Hla* RGH2). Branch support was generated using 1,000 bootstraps, with orange dots designating support from 80-100%. Species abbreviations listed in Supplementary Data 1. Rice (*Os*; *AK071926* (=*LOC_Os12g18360.2*)) used as an outgroup.

**Supplementary Fig. 4.**
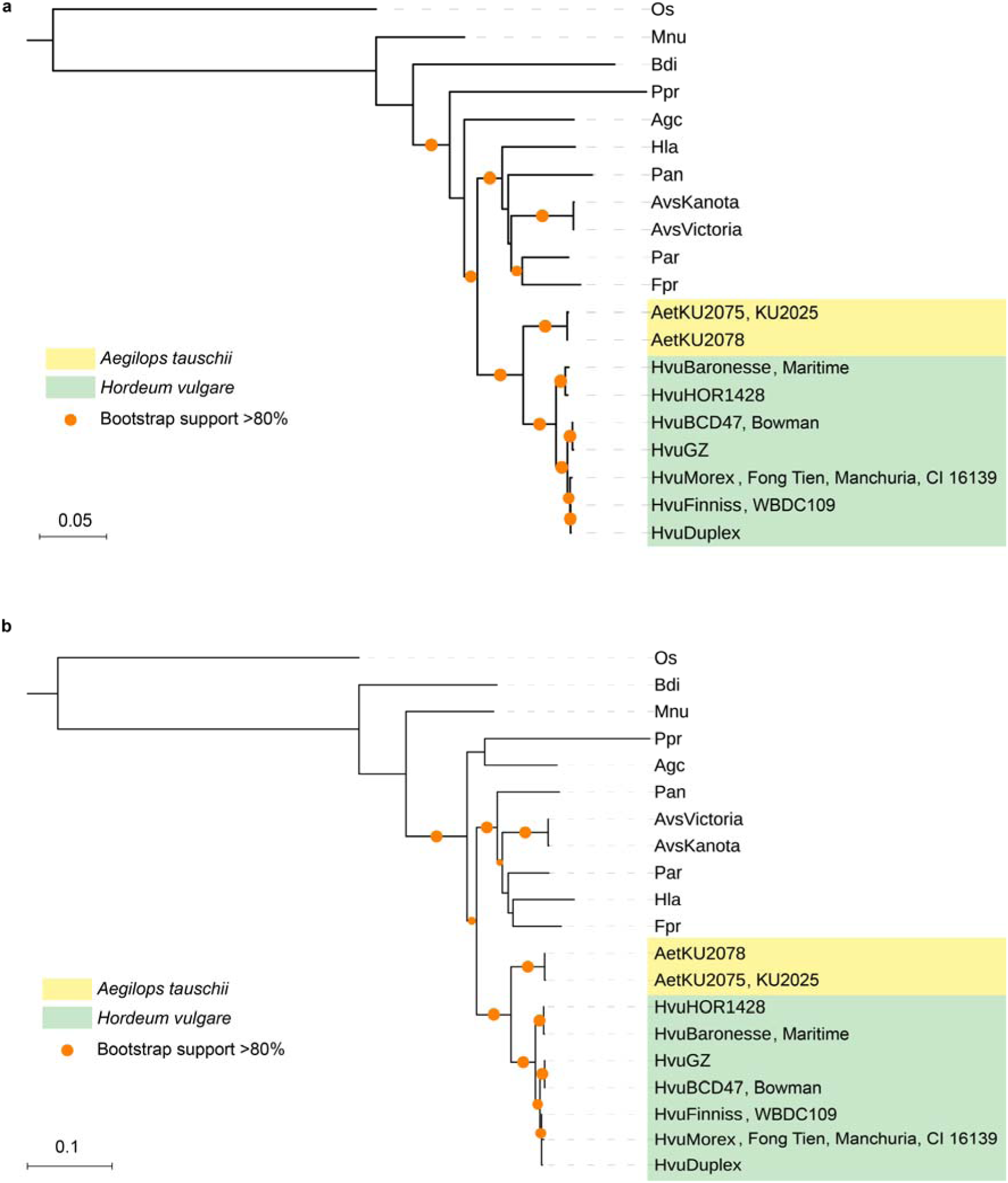
Maximum likelihood phylogenetic tree of *RGH3* homologs across the Poaceae. Alignments and phylogenetic trees of full length (a) and NB domain analyses (b) were performed. *RGH3* from *Ae. tauschii* and *H. vulgare* indicated with yellow and green highlighting respectively. Branch support was generated using 3,000 and 10,000 bootstraps for *RGH3* full length and NB domain, respectively, with orange dots designating support from 80–100%. Species abbreviations listed in Supplementary Data 1. Rice (*Os*; *LOC_Os12g18374.1*) used as an outgroup.

**Supplementary Fig. 5.**
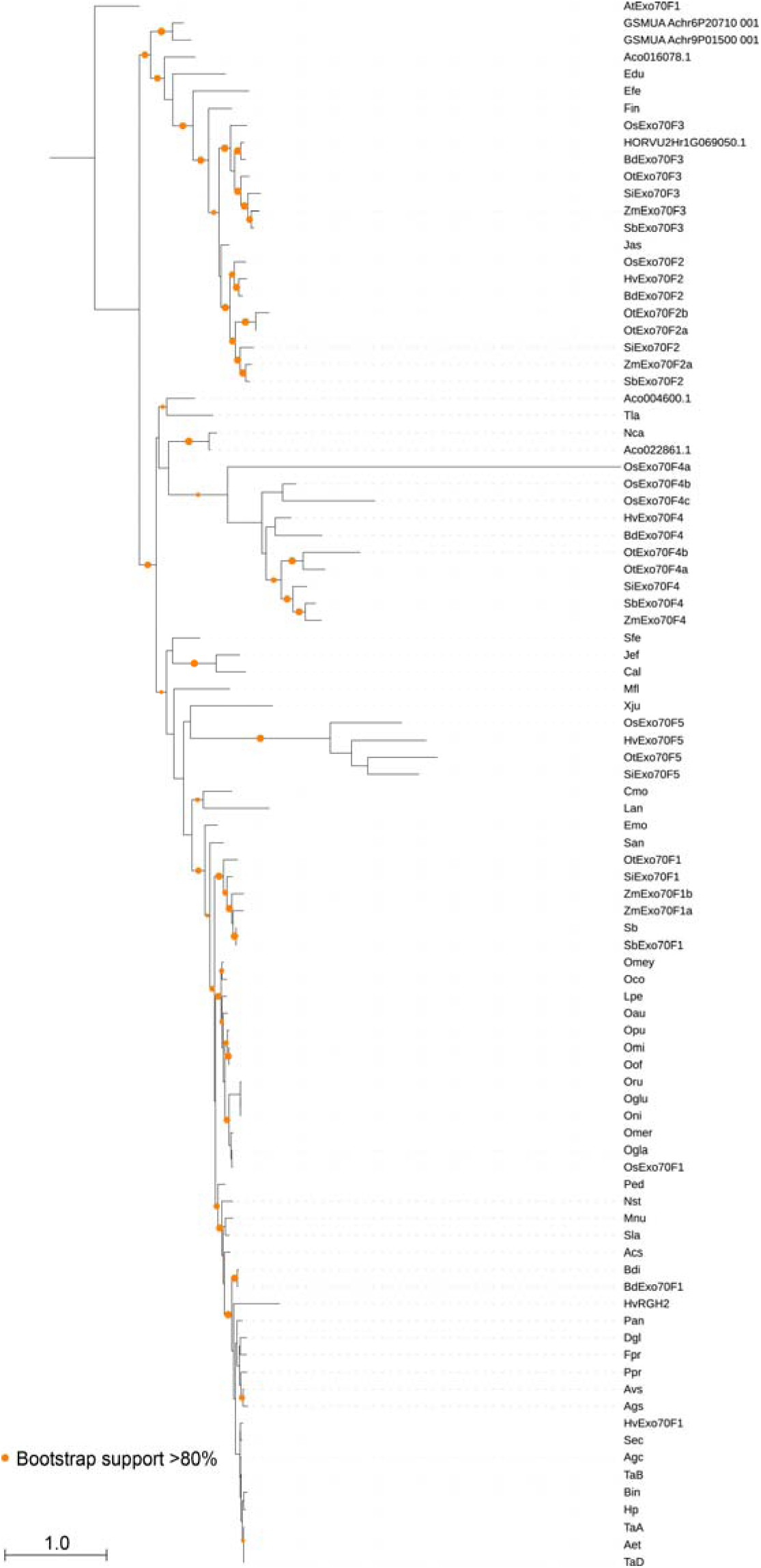
Maximum likelihood phylogenetic tree of *Exo70F* family members from diverse Poales species. Branch support was generated using 1,000 bootstraps, with orange dots designating support from 80–100%. Species abbreviations listed in Supplementary Data 1. *A*. *thaliana* (*At*; *AT5G50380.1*) used as an outgroup.

**Supplementary Table 1.**
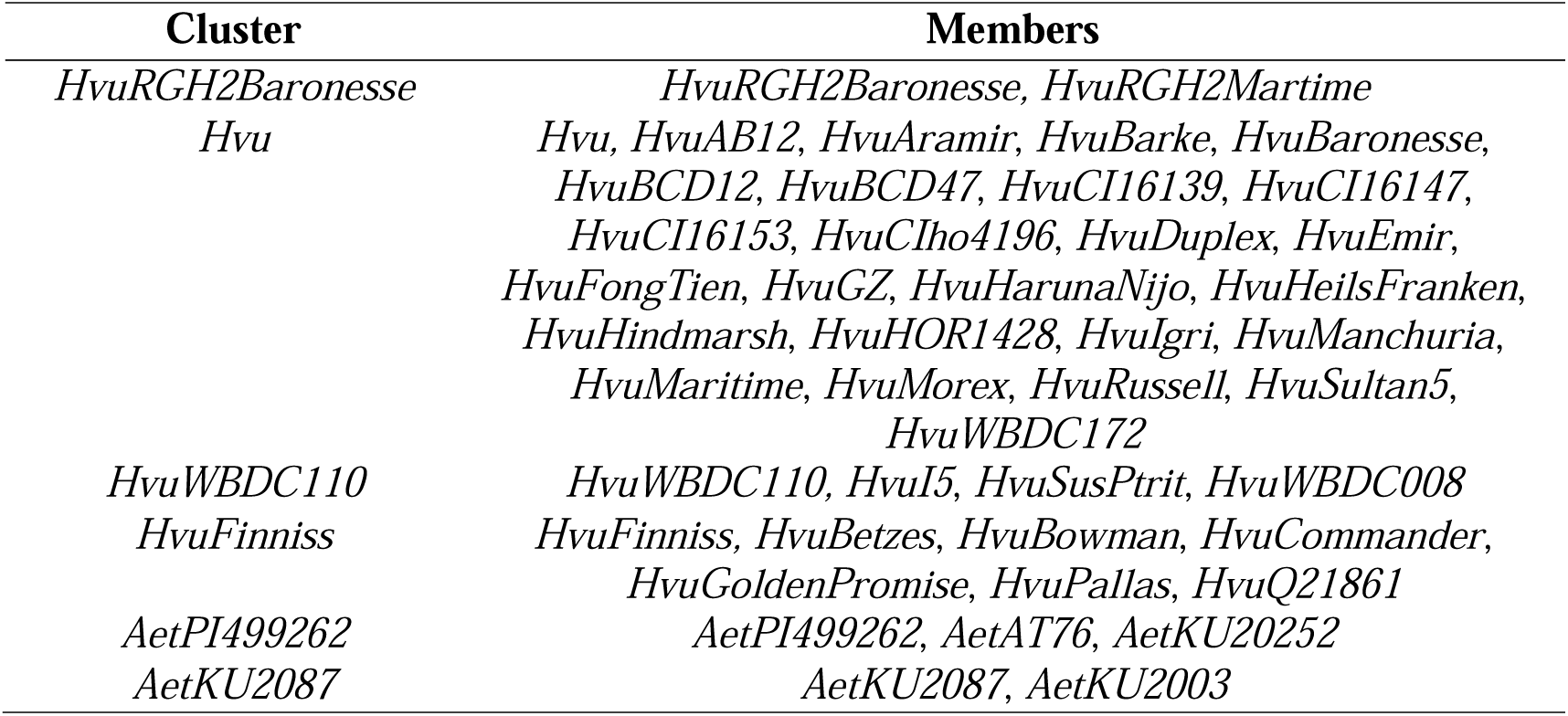
Identical gene clusters in *Exo70F1* phylogenetic analysis.

**Supplementary Table 2.**
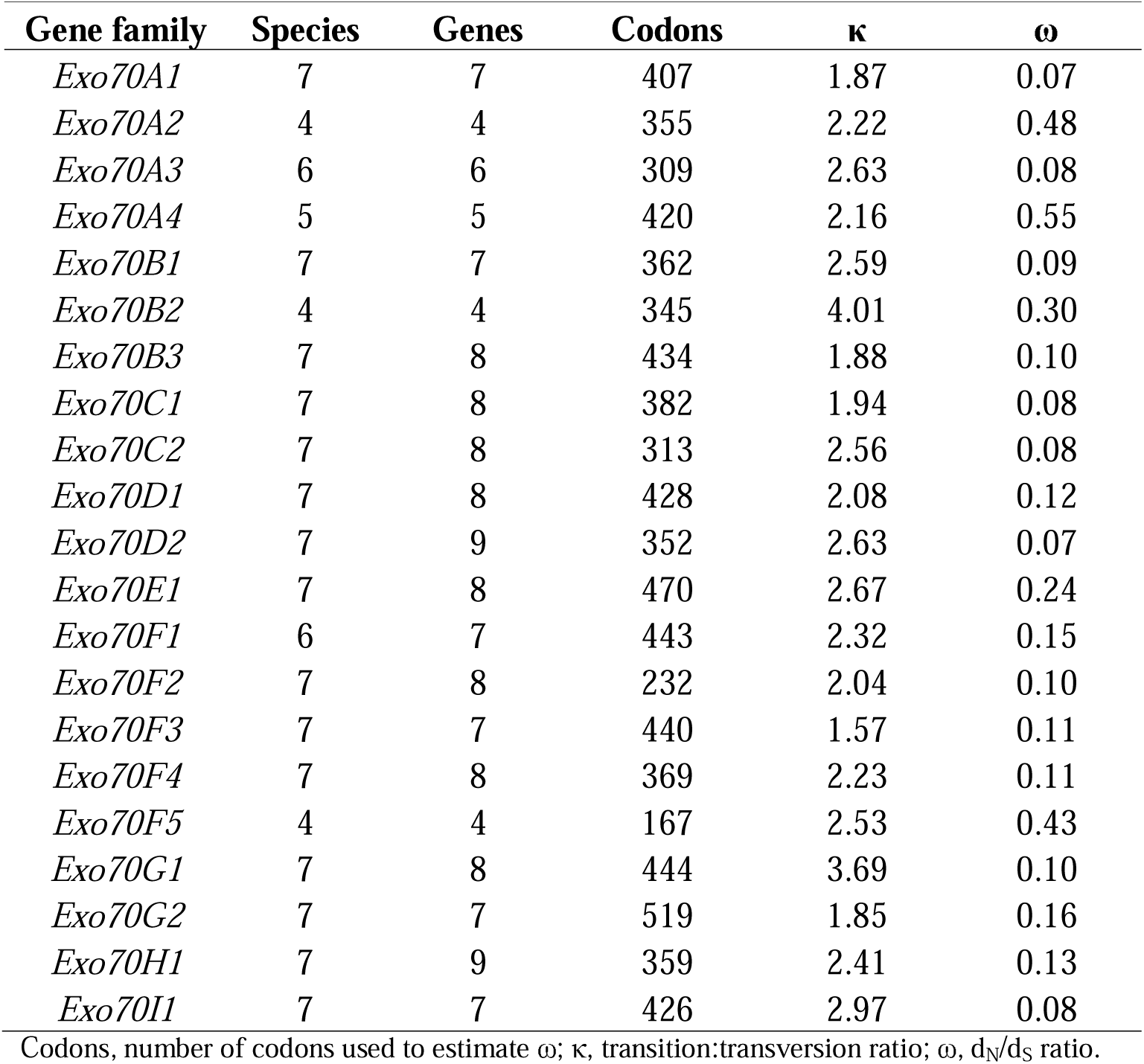
Parameter estimates of d_N_/d_S_ (ω) in different *Exo70* gene families.

**Supplementary Table 3.** Parameter and log likelihood estimates of d_N_/d_S_ (ω) in non-integrated and integrated *Exo70F1* homologs.

| Model | $\kappa$ | $\omega_0$ | $\omega_1$ | $\omega_a$ | $\omega_b$ | lnL |
| --- | --- | --- | --- | --- | --- | --- |
| H<sub>0</sub> (one $\omega$) | 2.403 | 0.213 | = $\omega_0$ | = $\omega_0$ | = $\omega_0$ | -11592.2 |
| H<sub>1</sub> (two $\omega$; $\omega_0$ and $\omega_1$) | 2.403 | 0.215 | 0.094 | = $\omega_0$ | = $\omega_0$ | -11591.3 |
| H<sub>2</sub> (two $\omega$; $\omega_0$ and $\omega_a$) | 2.422 | 0.102 | = $\omega_a$ | 0.423 | = $\omega_0$ | -11486.5 |
| H<sub>3</sub> (three $\omega$; $\omega_0$, $\omega_a$, $\omega_b$) | 2.419 | 0.078 | = $\omega_a$ | 0.423 | 0.132 | -11480.4 |

Significant differences in $\omega$ were observed for H<sub>0</sub> vs. H<sub>2</sub> ( $p < 0.001$ ) and H<sub>2</sub> vs. H<sub>3</sub> ( $p < 0.001$ ). Variables: $\kappa$ , transition:transversion ratio, $\omega$ $d_N/d_S$ ratio, lnL, log-likelihood. All likelihood ratio tests had degrees of freedom of 1.

**Supplementary Data 1**. Identifiers of species, accessions, genomes, transcriptomes, and genes

**Supplementary Data 2**. Identifiers of *RGH2* and *RGH3* genomic contigs

**Supplementary Data 3**. XML file containing NLR-associated MEME motifs

**Supplementary Data 4**. Gene identifiers for *Exo70* gene family

